# GLUT5 armouring enhances adoptive T cell therapy anti-tumour activity under glucose-limiting conditions

**DOI:** 10.1101/2025.01.16.633365

**Authors:** Robert Page, Olivier Martinez, Daniel Larcombe-Young, Eva Bugallo-Blanco, Sophie Papa, Esperanza Perucha

**Author notes:** Joint senior authors (Sophie Papa, Esperanza Perucha).

## Abstract

Cancer immunotherapy with engineered T cells has become a standard treatment for certain hematologic cancers. However, clinical trial outcomes for solid tumours are significantly lagging. A primary challenge in solid tumours is the lack of essential metabolites in the tumour microenvironment, such as glucose, due to poor vascularization and competition with tumour cells. To address this, we modified T cells to use fructose as an alternative energy source by introducing ectopic GLUT5 expression. We show that “GLUT5-armored” T cells, engineered with either chimeric antigen receptors (CARs) or an ectopic T cell receptor (TCR), achieve enhanced anti-tumour activity in low-glucose environments in both *in vitro* and *in vivo* models. This straightforward modification is compatible with current clinical approaches and may improve the efficacy of T cell therapies for solid tumours.

**Graphical Abstract:** 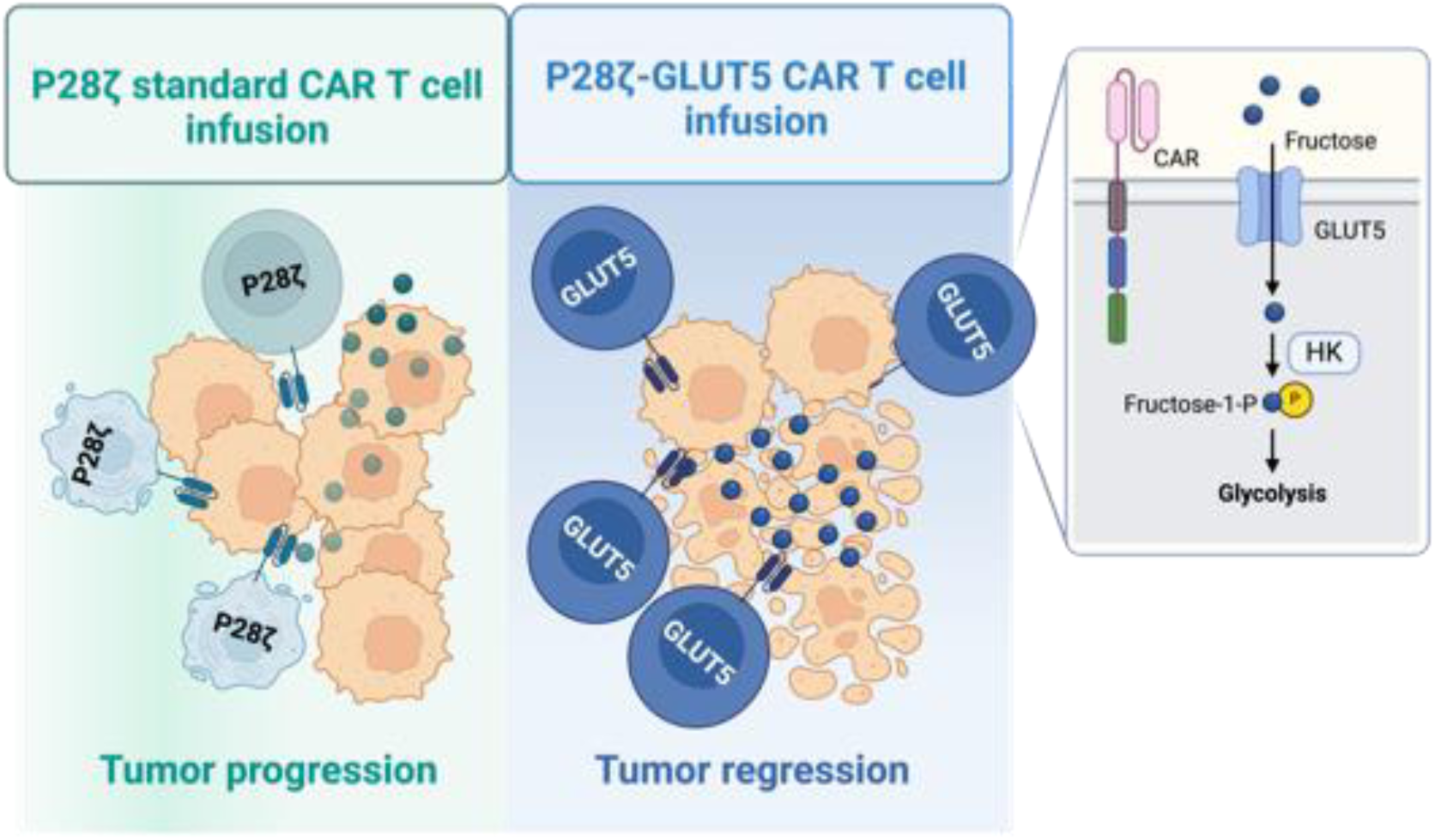

## Introduction

Adoptive cell therapy (ACT) has emerged as a cornerstone in cancer immunotherapy, showing promise in both hematologic and solid tumours, as well as applications in regenerative medicine and autoimmunity. In cancer treatment, ACT methods vary from minimally modified approaches, like tumour-infiltrating lymphocyte (TIL) and donor lymphocyte infusions, to advanced genetically engineered therapies that use selected T cell receptors (TCRs) or synthetic chimeric antigen receptors (CARs). These CAR T cell therapies have notably improved survival outcomes in advanced B cell malignancies by targeting markers like CD19^1–3^ and B cell maturation antigen (BCMA)^4,5^.

Despite successes in hematologic cancers, the efficacy of ACT in solid tumours remains suboptimal^6–8^. Clinical trials indicate that while CAR T cells can induce cytotoxicity in solid tumours, this effect is often limited to a minority of patients, highlighting the need for further optimization. One avenue for improvement is to engineer T cells for increased resilience against the hostile conditions of the solid tumour microenvironment (TME), which presents multiple barriers to ACT effectiveness^9^.

Resistance to ACT in solid tumours arises from both intrinsic and extrinsic factors. Intrinsic challenges include poor T cell homing, limited persistence post-transfer, and functional exhaustion from chronic stimulation. Extrinsic challenges, such as inhibitory signalling through immune checkpoints and suppressive cell types within the TME, further inhibit T cell activity^10,11^. Additionally, the TME’s altered metabolic environment—marked by limited glucose availability due to tumour cell competition—restricts T cell function^12^. Effector T cells rely on glycolysis even in the presence of oxygen^13–15^, a process essential for energy production and to generate metabolic intermediates essential for biosynthesis of proteins and nucleic acids, which is severely compromised in the hypoglycemic TME^16,17^.

To address this metabolic limitation, we propose equipping CAR and TCR T cells with the ability to metabolize fructose, a carbon source not typically utilized by most tumours, by ectopic GLUT5 expression. In this study, we demonstrate that fructose can serve as an alternative energy source, preserving T cell function under glucose-deficient conditions. We demonstrate that this metabolic flexibility enhances T cell efficacy in *in vivo* solid tumour environments, providing a new approach to improving ACT outcomes in solid cancers.

## Materials and Methods

### DNA

Sequences encoding 4ab, P28ζ, PTr or GLUT5 were cloned downstream of the human EF1a into the lentiviral transfer vector pUltra (Addgene plasmid # 24129). Construction of DNA vectors was conducted using NEBuilder HiFi DNA Assembly Kit as per manufacturer instructions (NEB). Assembly reactions were transformed by mixing the entire reaction with 25 mL of NEB 5-alpha competent E. coli (NEB) as per manufacturer instructions. Ampicillin resistant bacterial colonies were transferred to 15 mL of LB broth containing ampicillin (Sigma) (100 mg/mL) and grown at 37°C for 18 hr at 220 RPM. Plasmid DNA was extracted from bacterial cultures with E.Z.N.A Plasmid DNA Mini Kit II (Omega Bio-Tek) as per manufacturer instructions.

### Cell Lines

HEK-293T, PC3-LN3, PC3-LN3-PSMA and MCF7 cells were cultured in Dulbecco’s Modified Eagle Medium (DMEM) (Gibco) supplemented with 10% Foetal Bovine Serum (FBS) (Sigma). In experiments where media containing different concentrations of glucose or alterative to glucose are used, PC3-LN3-PSMA or MCF7 cells were cultured in glucose-free DMEM supplemented with 10% FBS. All cell lines were cultured at 37°C in 5% CO2.

### Lentiviral Production In HEK-293T

1×10^7^ HEK-293T cells were plated in 150 mm dishes and allowed to adhere for 24 hrs. HEK-293T cells were then transfected with 25 mg of lentiviral transfer plasmid, 12.5 mg pVSV-G and 12.5 mg of pCRV-1 (kind gifts from Dr Caroline Hull, KCL). Transfection reactions were prepared by mixing plasmids with 150 mg of polyethylenimine (Sigma) in 1 mL of serum-free DMEM and incubating at room temperature for 25 mins before reactions were added dropwise to cells. 24 hrs after transfection, media was aspirated and replaced with 17.5 mL of fresh media. 48 hrs after transfection, virus-containing media was removed and transferred to a 50 mL conical tube which was stored at 4°C. 17.5 mL of fresh media was added to cells. 72 hrs after transfection, virus-containing media was again removed from cells and pooled with the virus-containing media removed from cells the previous day. Lentivirus was then concentrated from virus-containing media by first centrifuging at 1500 g for 5 min to remove detached cells and debris. The supernatant containing lentivirus was transferred to a new 50 mL conical tube and centrifuged at 20000 g for 90 mins. The supernatant was aspirated, and the virus-containing pellet was resuspended in 200 mL of phosphate-buffered saline (PBS). Concentrated lentivirus was stored at -80°C until used.

### Isolation, culture, transduction and expansion of human PBMCs

PBMCs were purified by carefully layering whole blood over 15 mL of Ficoll-Paque (GR Healthcare), before centrifugation at 800 g for 35 min with brake and accelerator set to minimum. PBMCs were then transferred to a clean 50 mL tube and washed twice in PBS, before being resuspended in an appropriate volume of media.

PBMCs were cultured in RPMI 1640 (Gibco) supplemented with 5% human AB serum (Sigma). PBMCs were counted and diluted to 1×10^6^ cells/mL and activated with 5 μg/mL Phytohemagglutinin-L (Sigma) in a 96-well plate. 24 hrs after activation, cultures were supplemented with 100 U/mL recombinant human IL-2 (rhIL-2) (R&D Systems) and 50 μL of concentrated lentivirus was then added to cells and mixed by gently pipetting. For untransduced cells, 50 mL of PBS was added in place of lentivirus.

48 hrs after transduction, PBMCs were washed by adding 1 mL of PBS and centrifuged at 800 g for 5 mins. PBMCs were resuspended in 500 mL of fresh media supplemented with 100 U/mL rhIL-2. For PBMCs transduced with constructs containing the chimeric cytokine receptor 4ab, cultures were supplemented with 3 ng/mL recombinant human IL-4 (rhIL-4) instead of rhIL-2. Cells were then expanded to sufficient numbers by diluting with fresh media and supplementation of cultures with 100 U/mL rIL-2 or 3 ng/mL rIL-4 every 2–3 days.

For metabolite determination experiments, 0.5ml of PBMCs at 1×106 cells/mL were activated with PMA (50 ng/mL) and ionomycin (0.5 mg/mL) (both from Sigma) in 48-well tissue culture plates and incubated for the indicated period of time.

### Co-culture of tumour cells lines with PBMCs

Target cell lines were plated in either a 96-well, 24-well or 6-well tissue culture plate at a density of 1×10^5^ cells/cm2. Cells were allowed to adhere for 24 hrs. Expanded PBMC were then counted, washed in excess PBS to remove all cytokine and resuspended in fresh media. PBMC were then added to wells containing target-expressing cells at the indicated E:T ratio and incubated for the indicated time period.

In experiments where defined concentrations of glucose or fructose were used, target cell lines were first plated in standard media and allowed to adhere overnight. Before the addition of PBMC, media was aspirated from target cell line, wells were washed with excess PBS, and standard media was replaced with glucose-free media. PBMC were then counted, washed in excess media, resuspended in glucose-free media and added to target cell line cultures. Glucose or fructose (both from Sigma) dissolved in PBS was then added to cultures to achieve the indicated concentration.

### Cytotoxicity Assay

Cytotoxicity of CAR transduced cells toward target cell lines was determined using 3-(4,5-dimethylthiazol-2-yl)-2,5-diphenyltetrazolium bromide (MTT) assay. After co-culture of PBMC and target cell lines, the remaining media was completely aspirated and replaced with fresh DMEM supplemented with 10% FBS and 500 mg/mL MTT (Sigma). Plates were then incubated for 90 min at 37°C in 5% CO2. All media was then aspirated and formazan crystals resulting from remaining viable target cells were solubilised in Dimethyl sulfoxide (DMSO) (Sigma). The absorbance of this solution was measured at 590nm using a FLUOstar Omega plate reader. Cytotoxicity was determined relative to control wells where target cells had been cultured in the absence of PBMCs. For cultures containing defined concentrations of glucose or fructose, cytotoxicity was determined relative to target cells cultured in the same concentration of glucose or fructose. The viability of target cell cultures was calculated using the formula: % Viability = (A^Coculture^/A^Target Only^) x 100

### Cell Surface Staining

For staining of cells with unconjugated or fluorescently conjugated primary antibodies, cells were washed twice in 1 mL of staining buffer (PBS supplemented with 5% FBS) and then incubated with 100 μL of primary antibody at the indicated dilution for 30 min at 4°C. Cells were then washed in 1 mL of staining buffer. If using primary conjugated antibodies, cells were resuspended in 300 μL of staining buffer and analysed by flow cytometry. If stained with an unconjugated primary antibody, cells were incubated with 100 μL of fluorophore-conjugated secondary antibody at the indicated dilution for 30 min at 4°C. Cells were then washed in 1 mL of staining buffer, resuspended in 300 μL of staining buffer and analysed by ow cytometry. The following antibodies were used for cell surface protein expression; mouse anti-myc (9E10, 1:4, in-house production), mouse anti-human CD3 FITC (UCHT1, 1:40, Biolegend), mouse anti-human PD-1 PE (EH12.2H7, 1:40, Biolegend), mouse anti-human CD69 APC (FN50, 1:40, Biolegend), Goat anti-mouse PR (Poly4053, 1:100, Biolegend).

All samples for ow cytometry were acquired using either a BD LSRFortessa Flow Cytometer and BD FACSDiva software or a Beckman CytoFLEX and CytExpert software. Flow cytometry data were analysed using FlowJo software (BD).

### Assessment of cell viability by flow cytometry

Cellular viability was determined by staining with FITC-conjugated Annexin V (Biolegend) and Zombie Near Infared (NIR) Live/Dead (L/D) stain (Biolegend). Cells were first washed in PBS before being resuspended in 100 mL of Annexin V Binding Buffer (Biolegend) containing Annexin V (1:20 dilution) and Zombie NIR L/D (1:1000 dilution) and incubated for 30 mins at 4°C. Cells were then washed twice in Annexin V binding buffer before being resuspended in 300 mL of Annexin V staining buffer and analysed by flow cytometry.

### Intracellular cytokine staining (ICS)

6 hours before intracellular cytokine staining, 5 μg/mL of brefeldin A (Biolegend) was added to PBMC cultures. Cells were washed twice in staining buffer and, if required, stained with antibodies against cell surface antigens as described above. Cells were then fixed by resuspending them in 200 μL of 4% formalin and incubated at room temperature for 20 mins. Fixed cells were then washed once in 1 mL of staining buffer before being resuspended in 100 μL of permeabilisation (perm) buffer (1% Bovine Serum Albumin, 0.1% Saponin in PBS) and incubated at 4°C for 15 mins. Intracellular antibodies were then added directly to cells to achieve the final dilution indicated and cells were incubated at 4°C for 15 mins. Finally, cells were washed twice in 1 mL of perm buffer, resuspended in 300 μL of perm buffer and analysed by ow cytometry. The following antibodies were used for ICS; mouse anti-human TNFa PE (Mab11, 1:20, Biolegend), mouse anti-human IL-2 PerCP/Cy5.5 (MQ1-17H12, 1:20, Biolegend), mouse anti-human INFg APC (4S.B3, 1:20, Biolegend).

### Cell Enumeration with CountBright Beads

For counting cells with CountBright beads, cells were first washed twice in 1 mL of staining buffer. If necessary, cells were then stained for cell-surface protein expression as described above. Cells were then resuspended in 275 μL of staining buffer and 25 μL of CountBright Beads (Invitrogen) were added to each sample. Samples were then analysed by flow cytometry and the number of cells events and bead events tabulated. The cell concentration was then determined as per manufacturer instructions.

### Analysis of proliferation with CellTrace Far Red

For analysing PBMC proliferation, PBMCs were counted, washed in excess PBS and resuspended to a density of 1×106 cells/mL in PBS containing 1μM CellTrace Far Red (Invitrogen). PBMC suspensions were then incubated at 37°C for 20 mins. Cells were then washed in excess RPMI supplemented with 5% human serum, resuspended in an appropriate volume of media and either cocultured with target cell lines or stimulated with PMA and Ionomycin. 72–96 hrs after stimulation, cells were washed twice in 1 mL of staining buffer. If necessary, cells were then stained for cell-surface protein expression as described above. Cells were then resuspended in 300 μL of staining buffer and Cell Trace Far Red fluorescent signal was determined by ow cytometry.

### Western Blot

Protein lysates were generated from PBMCs by first resuspending cells in 100 μL of Radioimmunoprecipitation Assay buffer (150 mM NaCl, 1% Triton-X100, 0.5% Sodium Deoxycholate, 0.1% SDS, 50 mM Tris pH 8.0) and incubating on ice for 30 mins. Lysates were then cleared by centrifugation at 16000 g for 30 mins at 4°C. Cleared lysates were transferred to 1.5 mL conical tubes and stored at -20°C until use. Total protein concentration of lysates was determined using the BCA Protein Assay Kit (Thermo scientific), as per manufacturer instructions.

For polyacrylamide gel electrophoresis, 20 μg of protein was first mixed with 2X sample loading buffer (4% SDS, 10% 2-ME, 20% glycerol, 0.004% bromophenol blue, 125 mM Tris-HCl pH 6.8) and boiled at 95°C for 5 mins. Lysates were then separated on NuPAGE 4%-12% Bis-Tris Mini Protein Gels (Invitrogen) at 120 V for 90 mins before transfer to a PVDF membrane at 100 V for 90 mins at 4°C. Membranes were then blocked by incubating in blocking buffer (5% non-fat dried milk dissolved in PBS containing 0.1% Tween-20 (PBS-T)) for 1 hr at room temperature. Membranes were then incubated for 18 hrs in primary antibody diluted in blocking buffer at 4°C. Membranes were washed 3 times in PBS-T before being incubated with HRP-conjugated secondary antibody diluted in blocking buffer for 1 h. Membranes were then washed 3 times in PBS-T before Enhanced Chemiluminescent System (ECL) was used to image the signal with a Chemidoc XRS + imaging system (Bio-rad). The following antibodies were used for western blotting; mouse anti-human GLUT5 (E-2, 1:50, SCBT), rabbit anti-human b-actin (D6A8, 1:100, CST), goat anti-mouse HRP (1:1000, Invitrogen) goat anti-rabbit HRP (1:1000, Invitrogen).

### The Cancer Genome Atlas (TCGA) Analysis

TCGA expression data and plots were obtained using GEPIA2 web server Expression DIY tool.

### Metabolite Quantification

Glucose, Fructose and Lactate concentrations were quantified from sera or cell culture media using D-Glucose/D-Fructose Assay Kit or D-Lactate Kit (Megazyme). For glucose/fructose concentration determination, 1 μL of cell culture supernatant, sera or glucose/fructose standards was dispensed into a 96 well plate. 100 μL of water, 5 μL of solution I (buffer) and 5 μL of solution II (NADP+/ATP) was added to each well, and optical absorbance was measured at 340 nm (A1). 1 μL of solution III (HK/G6P-DH) was added to each well, samples were gently mixed, incubated at room temperature for 5 mins and optical absorbance was measured again at 340 nm (A2). Finally, 1 μL of solution IV (PGI) was added to each well and mixed gently. The plate was incubated at room temperature for 10 mins before optical absorbance was measured for a nal time at 340 nm (A3). Absorbance for glucose was calculated as A2-A1. Absorbance for fructose was calculated as A3-A2-A1. Absolute glucose and fructose concentrations were determined by plotting a standard curve of absorbance against the concentration of glucose/fructose standards, and interpolating the absorbance observed in each sample against this standard curve.

For lactate concentration determination, 1 μL of cell culture supernatant or lactate standards was dispensed into a 96 well plate. 100 μL of water, 5 μL of solution I (buffer), 5 μL of solution II (NADP+/PVP) and 1 μL of solution III (D-GPT) was added to each well, and optical absorbance was measured at 340 nm (A1). 1 μL of solution IV (LDH) was added to each well, samples were gently mixed, incubated at room temperature for 5 mins and optical absorbance was measured again at 340 nm (A2). Absorbance for lactate was calculated as A2-A1. Absolute lactate concentrations were determined by plotting a standard curve of absorbance against the concentration of lactate standards, and interpolating the absorbance observed in each sample against this standard curve. In all experiments absorbance was measured using a FLUOstar Omega plate reader.

### Xenograft Model and adoptive CAR T-cell therapy

NSG mice were maintained under specific-pathogen free conditions. The pharmacokinetics of intraperitoneal (IP) fructose delivery was determined by injecting mice with 300 mg/kg fructose dissolved in PBS via IP injection. At specified time points post-IP injection, 20 μL blood samples were collected from mice tail veins. Serum samples were obtained by centrifuging blood samples at 2000 g for 10 mins and transferring serum to a 1.5 mL conical tube. Samples were stored at -20°C until use. For CAR efficacy studies, mice were subcutaneously inoculated with 2.5×106 PC3-LN3-PSMA cells. 9 days after tumour engraftment, mice were intravenously injected with 1×106 CAR+ T cells. After CAR T cell adoptive transfer, mice received daily intraperitoneal injections of fructose (300 mg/kg). Tumour growth was monitored every 2–3 days by callipers measurement. Tumour volume was calculated as length × width^2^ × (π / 6). Mice weight was monitored every 7 days. For survival analysis, mice were humanely culled when either (1) tumour measured > 13.5mm in any direction, (2) tumour ulcerated, (3) > 15% weight loss or (4) mice exhibited poor mobility or piloerection.

### Statistics

Data were expressed as mean ± SEM. Statistical tests were performed using GraphPad Prism software (version 10, GraphPad). Differences between groups were analyzed for statistical significance by Student’s *t* test, or 2-way ANOVA as indicated.

### Study Approval

All animal experiments were conducted in accordance with UK Home Office guidelines under the project licence P23115EBF. Whole blood was obtained from healthy human donors, after written informed consent, under the approval of the Guy’s and St Thomas’ Research Ethics Committee (reference 09/H0804/92).

## Results

### Glucose availability is a limiting factor for CAR T cell function *in vitro*

To evaluate the effect of glucose availability on CAR T cell function, we used a prostate-specific membrane antigen (PSMA)-targeting CAR T cell model (P28ζ) known for strong *in vitro* performance but limited clinical efficacy^18^. This CAR consists of a PSMA-specific scFv joined to CD28 and CD3ζ signalling domains, allowing for efficient target recognition and activation. As a control, we used a truncated CAR (PTr) lacking intracellular signalling domains, designed to bind PSMA without activating T cell functions (Supplementary Figure 1A). As a CAR target cell, we used the cell line PC3-LN3-PSMA, derived from parental the PC3-LN3 line which have been stably transduced to overexpress PSMA.

Both P28ζ and PTr CAR constructs were effectively transduced into peripheral blood mononuclear cells (PBMCs) (Supplementary Figure 1B). We confirmed that PSMA expression was high in the PC3-LN3-PSMA cell line but absent in the parental PC3-LN3 line (Supplementary Figure 1C). Cytotoxicity assays showed that P28ζ CAR T cells effectively reduced the viability of PC3-LN3-PSMA cells, especially at high effector-to-target (E:T) ratios, whereas PTr cells showed minimal cytotoxicity, confirming specific CAR-mediated cytotoxic activity (Supplementary Figure 1D-E).

Next, we examined how different glucose concentrations affected P28ζ CAR T cell function. In co-culture experiments with a fixed 1:1 E:T ratio, we observed that P28ζ cytotoxicity was enhanced as glucose concentration increased up to 7 mM, reaching a plateau (Figure 1A). Similarly, CAR T cell proliferation followed this trend, with enhanced proliferative rates observed at higher glucose levels (Figure 1B). At low glucose concentrations (1 mM), mimicking the TME, the activation markers PD-1 and CD69 remained unchanged, indicating intact target recognition by the CAR T cells (Figure 1C). However, hypoglycemia markedly reduced the expression of interferon-gamma (IFNγ) and, to a lesser degree, tumour necrosis factor-alpha (TNFα), key indicators of effector function, while interleukin-2 (IL-2) production was unaffected (Supplementary Figure 2A). The reduction in these effector functions was confirmed to not be due to decreased CAR T cell viability at low glucose concentrations (Figure 1D).

**Figure 1.**
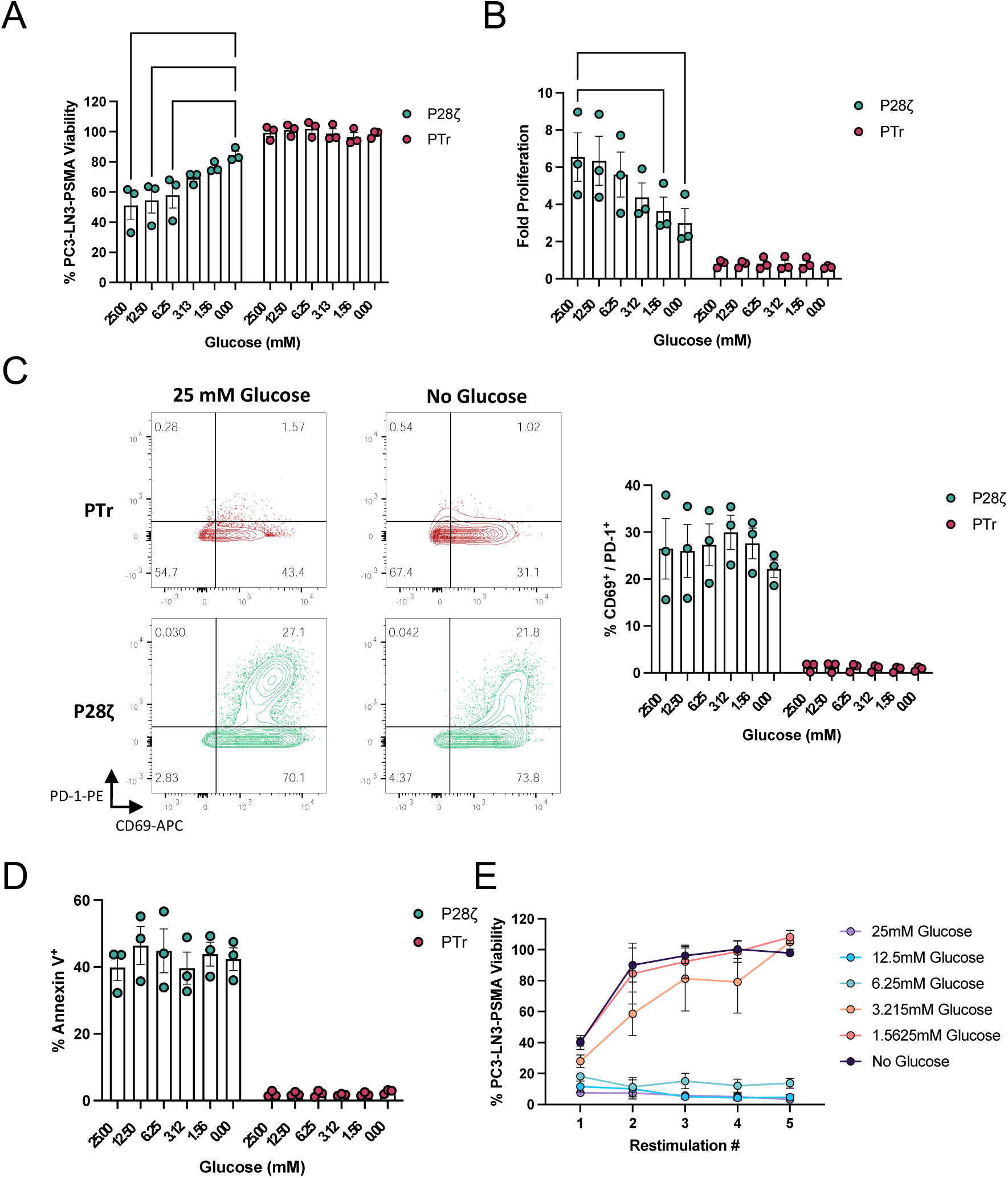
Glucose deprivation attenuates P28ζ CAR T cell function. **(a**) PC3-LN3-PSMA viability after 72 hrs co-culture with P28ζ or PTr CAR T cells in media supplemented with increasing concentrations of glucose. **b-d** (**b**) Fold proliferation; (**c**)CD69 and PD-1 expression; and (**d**)Annexin V binding of P28ζ or PTr CAR T cells co-cultured with PC3-LN3-PSMA cells for 72 hrs in media supplemented with increasing concentrations of glucose. **f.** PC3-LN3-PSMA viability after each restimulation round when co-cultured with serially restimulated P28ζ cells in the presence of increasing concentrations of glucose. Statistical significance relative to 25mM glucose conditions was calculated by two-way ANOVA. Data represents mean ± SEM of three independent healthy donors. * p≤0.05, ** p≤0.01 and *** p≤0.001

Further experiments showed that low glucose levels impaired the ability of CAR T cells to perform serial killing across multiple rounds of target cell re-challenge, as would be the case for CAR T cells encountering large numbers of transformed cells *in* vivo. When re-stimulated up to five times on PSMA-expressing targets, P28ζ cells maintained cytotoxicity only under high glucose conditions, whereas cytolytic activity was rapidly lost under low-glucose conditions (Figure 1E). These results collectively indicate that low glucose concentrations, akin to TME, significantly restricts CAR T cell cytotoxicity, proliferation, and cytokine production, although target-recognition and viability remain largely intact.

### Ectopic expression of GLUT5 enables fructose to fuel effector functions in PSMA-targeting CAR T cells

Given the metabolic competition within the TME, we aimed to engineer CAR T cells to use fructose as an alternative carbon source (ACS), circumventing their dependency on glucose. We chose to introduce GLUT5, a fructose-specific transporter, to allow CAR T cells to metabolize fructose, which is not ordinarily available to most tumours due to a lack of GLUT5 expression by transformed cell types (Supplementary Figure 3A). To assess the potential for selective fructose use, we queried the TCGA and GTEx RNA sequencing databases, confirming that GLUT5 expression is low across most tumour and tissue types (Supplementary Figure 4A). This observation suggested that other cell types in the TME would not likely compete for fructose, allowing armoured CAR T cells to benefit exclusively from this additional energy source. We also confirmed within our model system that the PC3-LN3-PSMA target cell line was unable to proliferate in fructose-supplemented media (Supplementary Figure 4B)

To evaluate our approach, we generated a modified CAR T cell construct by cloning the GLUT5 sequence upstream of the PSMA-targeting CAR (P28ζ) separated by a viral 2A skipping peptide for multi-cistronic expression, resulting in the GLUT5_4P28ζ CAR construct (Figure 2A). In addition, we incorporated a synthetic cytokine receptor, 4αβ (4ab), into the construct to enable selective T cell expansion in IL-4-supplemented media. This system allows for controlled growth of transduced T cells, a useful feature for CAR T cell production and scalability^19^. Control constructs included the 4P28ζ CAR (without GLUT5) and 4PTr, a truncated CAR that binds PSMA without activating T cell functions. We confirmed high CAR expression (>90%) in all constructs by flow cytometry (Figure 2B) and validated GLUT5 expression in GLUT5_4P28ζ cells through Western Blotting (Figure 2C).

**Figure 2.**
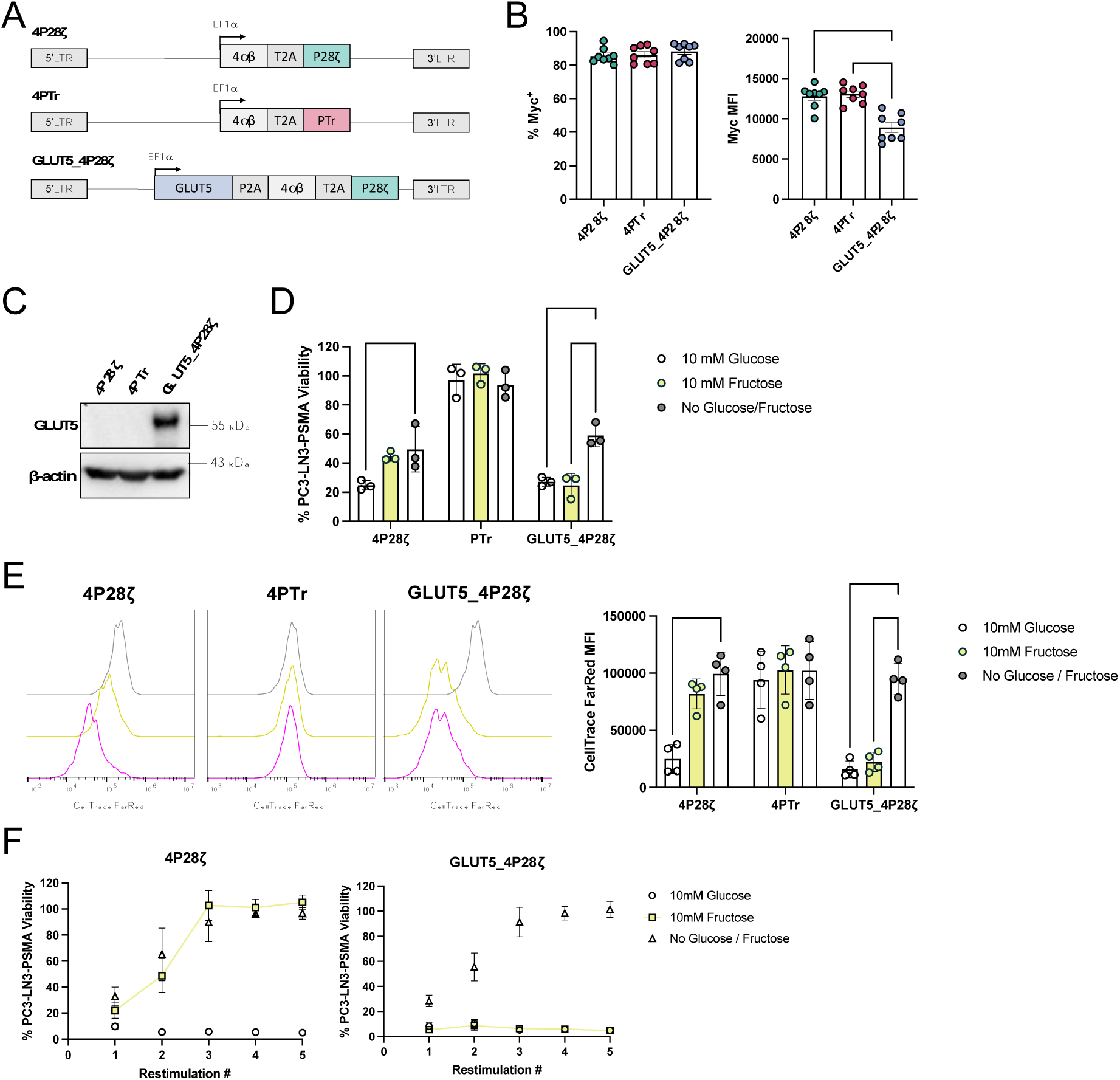
Ectopic expression of GLUT5 and media supplementation with fructose rescues 4P28ζ cytotoxicity and proliferation in the absence of glucose. **a.** 4P28ζ, 4PTr and GLUT5_4P28ζ construct design. **b.** 4P28ζ, 4PTr and 4P28ζ_GLUT5 transduction efficiency and CAR mean fluoresce intensity (MFI) as defined by staining for the CAR Myc tag. Statistical significance was calculated by two-way ANOVA. **c.** Representative GLUT5 and β-actin expression by 4P28ζ, 4PTr and GLUT5_4P28ζ as determined by western blotting. **d.** PC3-LN3-PSMA viability after co-culture with 4P28ζ, 4PTr or 4P28ζ_GLUT5 cells for 72 hrs in media containing no glucose/fructose, 10 mM glucose or 10 mM fructose. **e.** Relative proliferation of 4P28ζ, 4PTr or 4P28ζ_GLUT5 cells after co-culture with PC3-LN3-PSMA cells for 72 hrs in media containing no glucose, 10 mM glucose or 10 mM fructose. **f.** PC3-LN3-PSMA viability after each restimulation round when co-cultured with serially-restimulated 4P28ζ or 4P28ζ_GLUT5 cells in media no glucose, 10 mM glucose or 10 mM fructose. Statistical significance relative to No glucose/fructose conditions was calculated by two-way ANOVA. Data represents mean ± SEM of three to seven independent healthy donors. ** p≤0.01, *** p≤0.001, **** p≤0.0001.

In cytotoxicity assays, we tested the killing efficiency of GLUT5_4P28ζ cells against PSMA-expressing targets in media supplemented with 10mM glucose, 10mM fructose, or no additional monosaccharide. As expected, 4P28ζ CAR T cells exhibited robust cytotoxicity in glucose-rich conditions but showed limited effectiveness in fructose-only media. In contrast, GLUT5_4P28ζ cells displayed strong cytotoxicity in both glucose and fructose media, demonstrating that GLUT5 enables fructose-driven cytotoxic activity (Figure 2D).

Next, we evaluated whether fructose could support T cell proliferation. We found that 4P28ζ cells proliferated only in glucose-supplemented media, whereas GLUT5_4P28ζ cells showed significant proliferation in both glucose and fructose (Figure 2E). These results confirm that GLUT5 expression enables CAR T cells to use fructose as a carbon source for cell division.

We then tested the functionality of GLUT5_4P28ζ cells under repeated antigen stimulation, simulating chronic engagement in the TME. While 4P28ζ cells lost cytotoxic capacity after multiple rounds of stimulation in fructose-only conditions, GLUT5_4P28ζ cells retained high levels of cytotoxicity in both fructose and glucose media (Figure 2F). This supports our hypothesis that GLUT5 expression allows CAR T cells to maintain effector functions even in glucose-limited environments, representing a novel approach to improving CAR T cell persistence and efficacy in solid tumours.

We next aimed to confirm that fructose could serve as an ACS for anaerobic glycolysis, which is essential for effector T cell function, in GLUT5-expressing CAR T cells. To do this, we activated PBMCs transduced with 4P28ζ, 4PTr, or GLUT5_4P28ζ constructs using phorbol 12-myristate 13-acetate and ionomycin (PMA+I), a method that stimulates T cells independently of target cell engagement. This approach eliminated the potential confounding effects of a target cell line, ensuring that any observed changes in metabolite levels could be attributed solely to CAR T cell activity.

We stained the transduced PBMCs with Cell Trace Far Red and activated them with PMA+I in media containing 10 mM glucose, 10 mM fructose, or no added monosaccharide. After 72 hours, we measured Cell Trace Far Red fluorescence to assess proliferation. The proliferation patterns following PMA+I stimulation mirrored those observed during PC3-LN3-PSMA target cell co-culture (Figure 3A). Specifically, 4P28ζ and 4PTr cells exhibited significantly greater proliferation in glucose-supplemented media compared to fructose or no-sugar conditions. In contrast, GLUT5_4P28ζ cells proliferated equally well in both glucose and fructose media, indicating that GLUT5 facilitates fructose-driven cell division in response to activation by either CAR-antigen or PMA+I.

**Figure 3.**
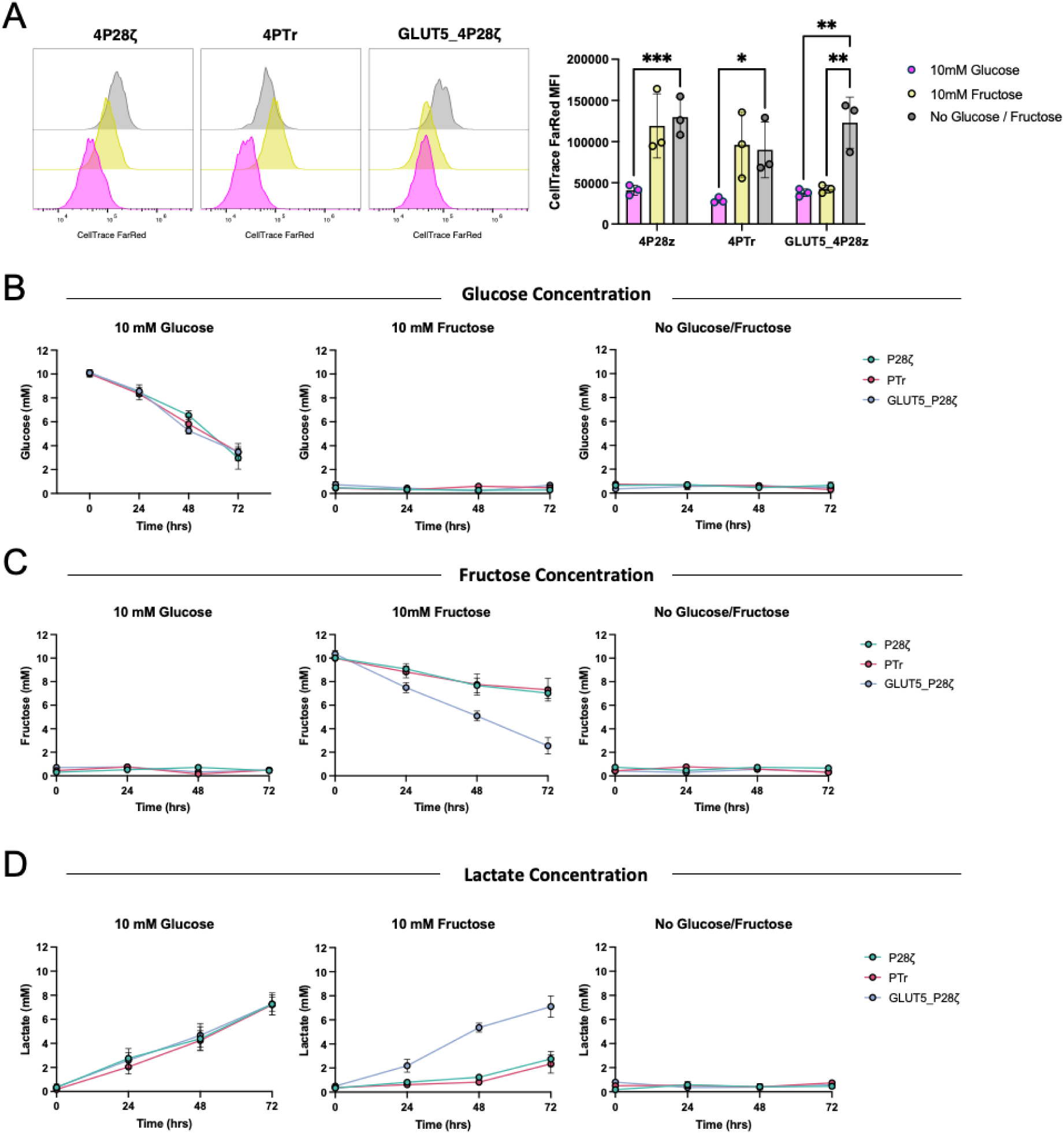
Ectopic GLUT5 expression enables fructose consumption and lactate production by activated T cells. **a.** Relative proliferation of 4P28ζ, 4PTr or GLUT5_4P28ζ cells after stimulation with PMA and Ionomycin for 72 hrs in media containing no glucose, 10 mM glucose or 10 mM fructose. (**b-d**) Cell culture supernatant from the experiment described in (a) was collected at 0, 24, 48 and 72 hrs and analysed for (b) glucose, (c) fructose and (d) lactate concentrations. Statistical significance relative to no glucose/fructose conditions was calculated by two-way ANOVA. Data represents mean ± SEM of three independent healthy donors. * p≤0.05, ** p≤0.01 and *** p≤0.001

To characterize GLUT5-mediated metabolism, we measured glucose, fructose, and lactate concentrations in the cell culture supernatants every 24 hours post-activation (Figure 3B-D). For all CAR constructs, glucose levels decreased linearly in glucose-supplemented media, accompanied by a reciprocal increase in lactate, indicating active anaerobic glycolysis. As expected, no fructose was detected in glucose-only media. However, in fructose-supplemented conditions, GLUT5_4P28ζ cultures rapidly depleted fructose with a corresponding increase in lactate, whereas fructose and lactate levels remained largely unchanged in 4P28ζ and 4PTr cultures. These findings indicate that ectopic GLUT5 expression enables CAR T cells to use fructose as an ACS for anaerobic glycolysis, resulting in lactate production and supporting proliferation and effector function in fructose-only environments.

### Ectopic expression of GLUT5 enhances anti-xenograft activity of PSMA-targeting CAR T Cells in response to intra-peritoneal fructose

After confirming that GLUT5_4P28ζ CAR T cells could proliferate and exert cytotoxic functions when supplied with fructose *in vitro*, we investigated their anti-tumour efficacy *in vivo*. For the ACT product expressing GLUT5 to effectively utilize fructose for its effector functions, sufficient fructose levels must be present in the serum and TME. This may necessitate oral or parenteral supplementation.

Initially, we assessed whether fructose could be delivered orally to mice. We supplemented the drinking water of NOD scid gamma (NSG) mice with 5% fructose for 72 hours. Serum samples were then collected to quantify glucose and fructose levels, comparing them to mice consuming standard drinking water. However, supplementation with 5% fructose did not result in a detectable increase in serum fructose concentrations. As an alternative, we administered fructose (300 mg/kg) through intraperitoneal (I.P.) injection and monitored serum fructose levels over 120 minutes post-injection. We detected basal serum concentrations of approximately 2 mM that rose rapidly, and returned to baseline levels by 30 minutes (Supplementary Figure 5A). These results indicate that there is a baseline fructose level in mice that may sustain GLUT5-expressing CAR T cells, and that I.P. administration of fructose temporarily elevates serum concentrations.

To evaluate the impact of GLUT5 expression on CAR T cell function *in vivo*, we established PC3-LN3-PSMA xenografts in NSG mice for nine days prior to administering 1 × 10^6^ transduced CAR T cells (4P28ζ, 4PTr, or GLUT5_4P28ζ) or PBS as a control via intravenous (I.V.) injection. We delivered 300 mg/kg of fructose daily through I.P. injection (Figure 4A). The 4P28ζ CAR T cells delayed xenograft progression compared to PBS or PTr-treated mice, consistent with previous reports. GLUT5_4P28ζ CAR T cells led to a significant further delay in xenograft progression and prolonged mouse survival compared to all other conditions in this aggressive tumour model (Figure 4B-C). Importantly, treatment with both CAR T cells and fructose was non-toxic, with no significant weight loss observed in any of the mice (Figure 4D). These findings underscore the potential of GLUT5_4P28ζ CAR T cells to enhance anti-tumour responses *in vivo*.

**Figure 4.**
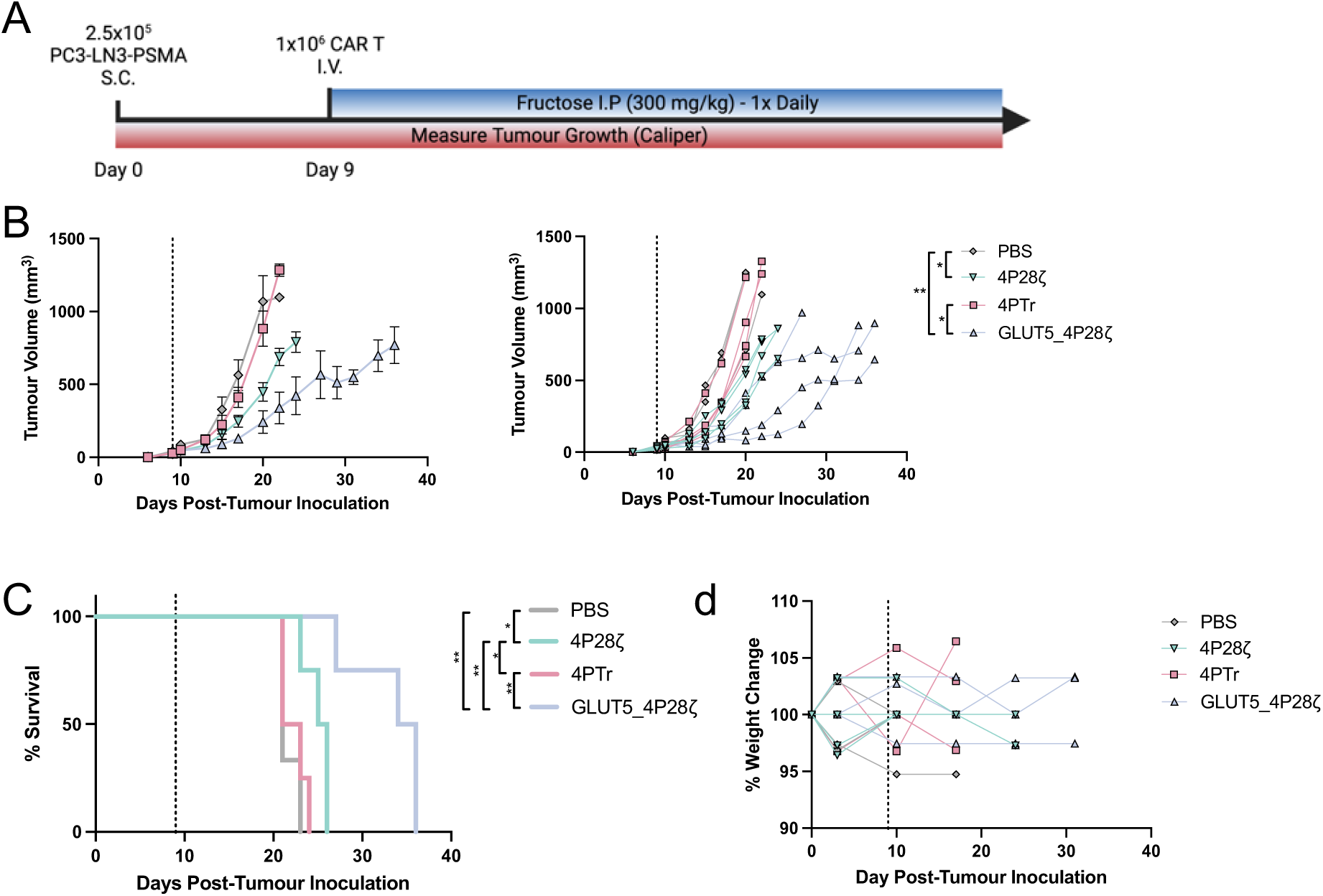
Fructose acts as an ACS for GLUT5 expressing CAR T cells *in vivo*. **a.** *in vivo*. tumour challenge experimental design. **b-d** (b) tumour growth for grouped and individual animals; (c) survival; and (d) weight change for mice inoculated with PSMA-LN3-PSMA cells and treated with either PBS (n=2), 4P28ζ (n=4), 4PTr (n=4) or GLUT5_ 4P28ζ (n=4) while receiving daily intraperitoneal (IP) fructose injections. Dashed line indicates time-point of PBS or CAR T cell infusion. Statistical significance of tumour volume was calculated by two-way ANOVA, statistical significance of mice survival was calculated by logrank test. * p≤0.05 and ** p≤0.01

### Ectopic expression of GLUT5 enables fructose to fuel effector functions of CAR or TCR engineered T Cell against diverse targets

To explore the versatility of fructose utilization by GLUT5-expressing T cells, we assessed the functional activity of CAR T cells engineered to target the epidermal growth factor receptor family member HER-2. We designed a HER-2-specific CAR construct by replacing the PSMA-specific scFv in our previously described constructs with an scFv derived from the monoclonal antibody trastuzumab, resulting in the derivatives 4H28ζ, 4HTr, and GLUT5_4H28ζ (Figure 5A). For our target cell line, we chose MCF7, a breast cancer cell line known for its HER-2 expression. In co-culture experiments, 4H28ζ CAR T cells exhibited enhanced cytotoxicity and proliferation with 10 mM glucose compared to conditions without glucose or fructose, but there was no improvement with 10 mM fructose. In contrast, GLUT5_4H28ζ CAR T cells demonstrated increased cytotoxicity and proliferation in both 10 mM glucose and 10 mM fructose, indicating their capacity to utilize fructose as an energy source for effector functions (Figure 5B-C).

**Figure 5.**
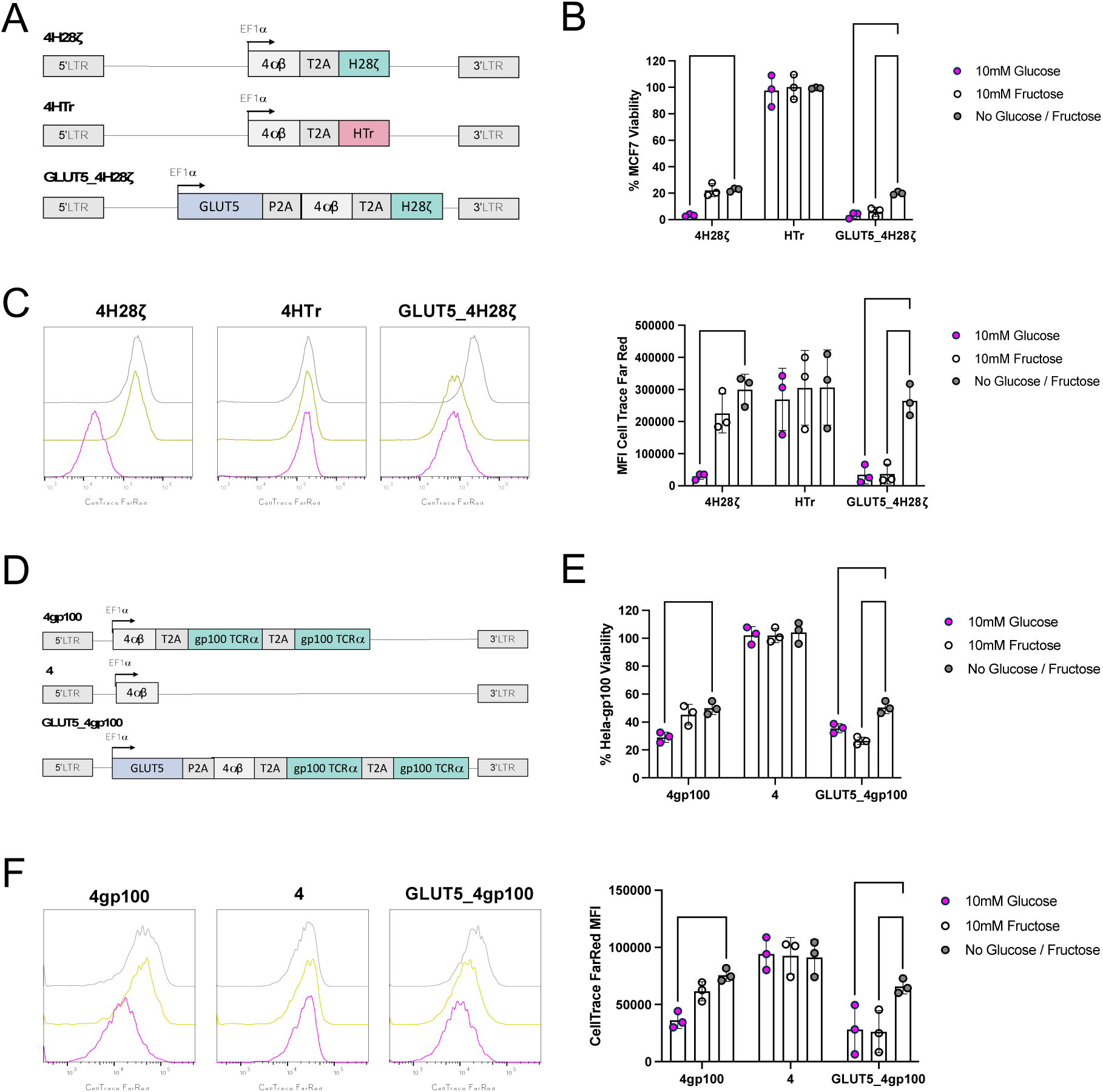
Fructose can be utilized as ACS by CAR T cells targeting HER-2 and by TCR-T cells targeting gp100. **a.** 4H28ζ, 4HTr and 4H28ζ_GLUT5 construct design. **b**. MCF7 viability after co-culture with 4H28ζ, 4HTr or GLUT5_4H28ζ cells for 72 hrs in media containing no glucose/fructose, 10 mM glucose or 10 mM fructose. **c.** Relative proliferation of 4H28ζ, 4HTr or GLUT5_4H28ζ cells after co-culture with PC3-LN3-PSMA cells for 72 hrs in media containing no glucose/fructose, 10 mM glucose or 10 mM fructose. **d.** 4gp100, 4 and 4gp100_GLUT5 construct design. **e.** Hela-gp100 viability after co-culture with 4gp100, 4 or 4gp100_GLUT5 cells for 72 hrs in media containing no glucose, 10 mM glucose or 10 mM fructose. **f.** Relative proliferation of 4gp100, 4 or 4gp100_GLUT5 cells after co-culture with Hela-gp100 cells cells for 72 hrs in media containing no glucose, 10 mM glucose or 10 mM fructose. Statistical significance relative to no glucose/fructose conditions was calculated by two-way ANOVA. Data represents mean ± SEM of three independent healthy donors. ** p≤0.01, *** p≤0.001, **** p≤0.0001.

Beyond CAR T cells, there is considerable interest in TCR-modified T cell therapies that target intracellular antigens presented as peptides on human leukocyte antigens (HLA). To investigate whether fructose utilization could benefit both CAR and TCR-engineered T cells, we expressed a gp100 peptide-specific TCR alongside either the 4αβ construct or with GLUT5 (Figure 5D). We conducted co-culture experiments with a gp100-expressing target cell line under conditions consistent with those used for the PSMA and HER-2-directed CAR T cell assays. Similar to the CAR T cell data, T cells engineered with the gp100 TCR showed significantly greater cytotoxicity and proliferation in the presence of 10 mM glucose compared to no glucose or fructose. However, the gp100-engineered T cells that also expressed GLUT5 exhibited significantly enhanced cytotoxicity and proliferation in both 10 mM glucose and 10 mM fructose (Figure 5E-F). These findings confirm that ectopic GLUT5 expression allows CAR and TCR T cells targeting a diverse range of antigens to effectively utilize fructose to support their effector functions.

## Discussion

In this study, we identified that glucose availability significantly limits the effector functions of CAR T cells, likely limiting their activity within the hypoglycaemic TME characteristic of solid tumours. This deficiency is likely to hinder the efficacy of CAR T cell therapies in combating solid malignancies^14,17^. To address this challenge, we hypothesized that engineering CAR T cells to utilize fructose through ectopic expression of GLUT5 could provide a simple yet effective solution. Our results demonstrate that ectopic GLUT5 expression enhances the ability of CAR T cells to utilize fructose as an alternative carbon source, effectively rescuing their effector functions *in vitro*. Metabolite analysis revealed that GLUT5-expressing CAR T cells can leverage fructose to fuel anaerobic glycolysis, leading to improved proliferation and cytotoxicity. This finding highlights the metabolic adaptability of CAR T cells when provided with access to alternative substrates.

In our *in vivo* studies, we established that GLUT5-expressing CAR T cells significantly delayed tumour progression and improved survival in xenograft models, further underscoring the potential of fructose utilization in enhancing CAR T cell efficacy. Additionally, we explored the application of this metabolic strategy in TCR-engineered T cells targeting the gp100 antigen. Similar to our findings with the HER-2 targeted CAR T cells, GLUT5-expressing TCR-modified T cells demonstrated increased cytotoxicity and proliferation in the presence of both glucose and fructose. These results collectively indicate that the approach of engineering T cells to utilize fructose is broadly applicable across diverse T cell engineering strategies.

While our study provides valuable insights into the potential use of fructose in enhancing CAR T cell efficacy against solid tumours, there are notable limitations to consider. A major limitation is that we cannot ensure if fructose exclusively fuels anaerobic glycolysis or if it is also shuttled to alternative metabolic pathways distinct from those utilized by glucose, because we did not directly trace intracellular fructose fate. Future studies employing radio-labelled fructose tracing could address this gap. Nonetheless, our results indicate that fructose utilization is not toxic and it does not negatively impact T cell effector functions, which is crucial for effective anti-tumour responses in clinical settings.

In our *in vivo* studies, we chose to supplement endogenous fructose levels exogenously via intraperitoneal (I.P.) injection. While this approach enhances fructose availability for GLUT5-expressing CAR T cells, it limits our ability to determine whether the observed functional benefits could be achieved solely through reliance on endogenous fructose concentrations. From a clinical perspective, while chronic metabolic conditions are associated with elevated long-term serum fructose levels^23^, short-term fructose supplementation in cancer patients would likely be justified if enhanced anti-tumour activity is observed in clinical trials. Thus, any potential requirement for exogenous fructose does not impede the clinical translation of our findings.

Our findings demonstrate that GLUT5 can be seamlessly incorporated into ACT genetic engineering constructs through the use of a 2A skip peptide, thereby avoiding any additional complexity in the manufacturing processes of these therapies. This simplicity in implementation enhances the feasibility of integrating metabolic engineering strategies into existing CAR T cell therapies. Previous reports have explored the engineering of cell lines^20^ and T cells^21,22^ to utilize fructose; however, this is the first study to directly examine such a system within the context of human ACT for solid tumours—a significant area of unmet clinical need where ACT has struggled to advance in clinical trials. This novelty underscores the relevance of our findings and emphasizes the need for further investigation into the clinical applicability of fructose-utilizing CAR T cells.

Moreover, our approach serves as a proof of concept: that engineering metabolic flexibility in T cells— which could be achieved not only through ectopic expression of transporters like GLUT5 but also possibly through the expression of diverse intracellular enzymes and metabolic regulators—represents a promising strategy to overcome the metabolic constraints imposed by the TME. By expanding the metabolic repertoire of T cells, we open new avenues for enhancing the efficacy of ACT in solid tumours, addressing a critical challenge in the field and paving the way for future clinical applications.

## Author Contributions

RP, SP & EP were responsible for Conceptualisation, Methodology and Writing – original draft. RP, OM, EBB & DLY were responsible for Investigation & Formal Analysis. OM, EBB and DLY were additional responsible for Writing – review & editing. SP and EP were responsible for Supervision.

## Acknowledgments & Funding

This research was supported by the National Institute for Health Research (NIHR) Biomedical Research Centre based at Guy’s and St. Thomas’ NHS Foundation Trust and King’s College London (RP). EP acknowledges funding from the Academy of Medical Sciences (SBF007\100069).

