## Supplementary figures for "GLUT5 armouring enhances adoptive T cell therapy anti-tumour activity under glucose-limiting conditions"

### Supplementary Material

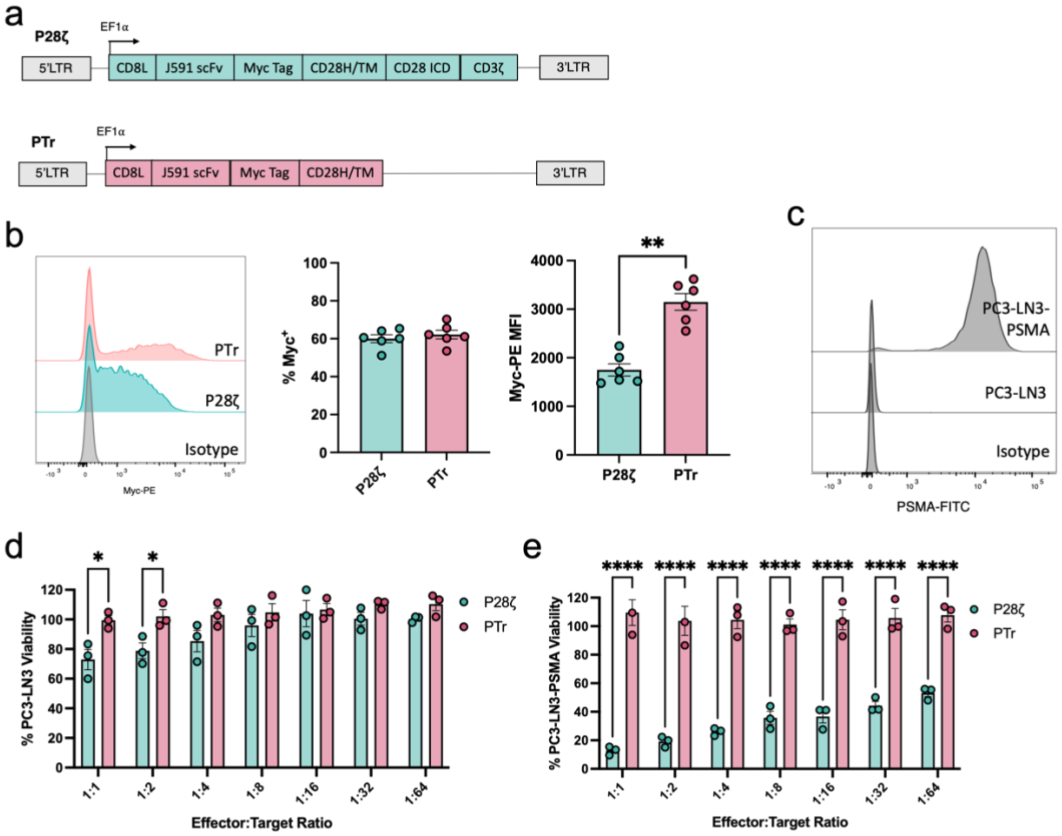


Supplementary Figure 1| **Anti-tumour activity of P28ζ CAR T cells. a**. P28ζ and PTr construct design. **b.** P28ζ and PTr transduction efficiency and CAR mean fluoresce intensity (MFI) as defined by staining for the CAR Myc tag. **c.** Representative surface PSMA expression by PC3-LN3 cells and PC3-LN3-PSMA cells. **d-e** (d) PC3-LN3 and (e) PC3-LN3-PSMA viability after 72 hr co-culture with either P28ζ or PTr cells at reducing effector:target ratios. Statistical significance calculated by two-way ANOVA. Data represents mean ± SEM of three to six independent healthy donors. * p≤0.05, ** p≤0.01, **** p≤0.0001.

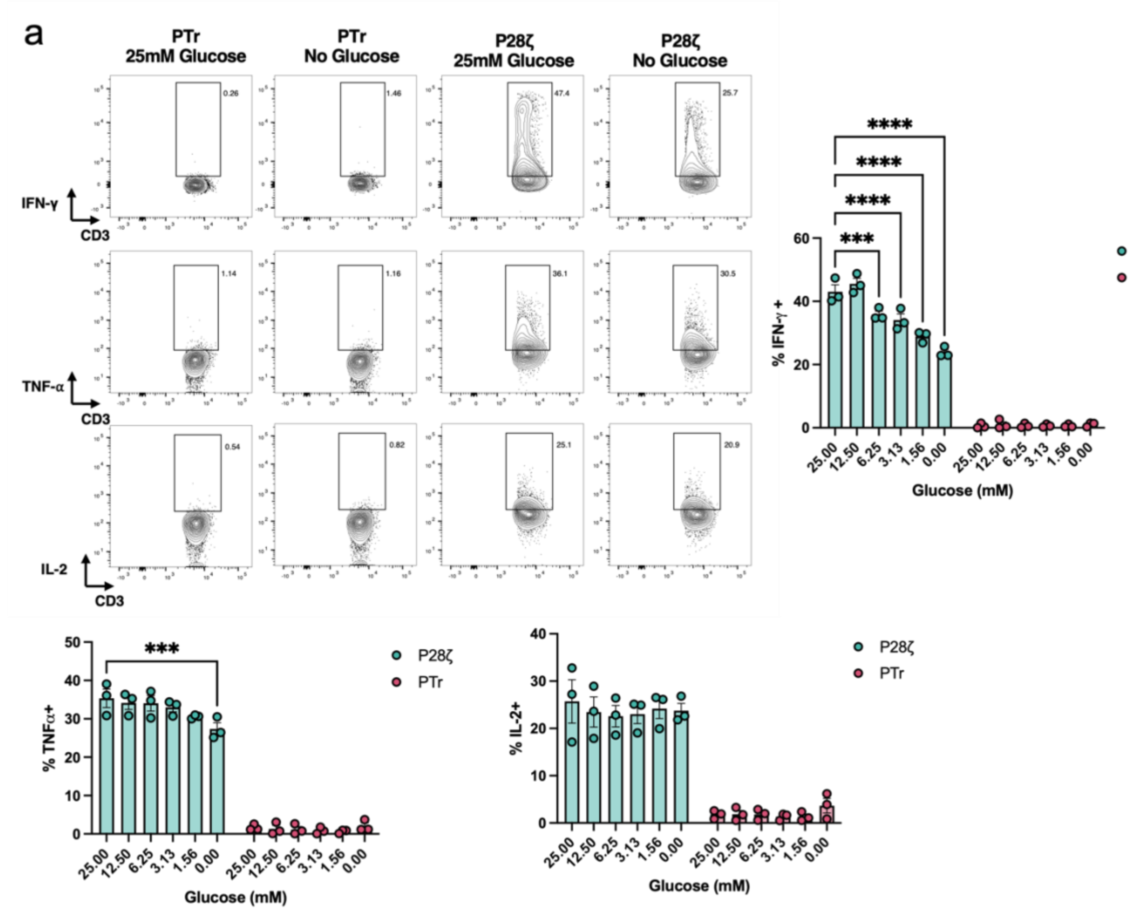


Supplementary Figure 2 | **Glucose availability has differential effects on pro-inflammatory cytokine production by P28ζ CAR T cell. a**. Intracellular IFNɣ, TNFα and IL-2 staining in P28ζ or PTr cocultured with PC3-LN3-PSMA cells for 72hrs in media containing increasing concentrations of glucose. Statistical significance relative to 25mM glucose conditions was calculated by two-way ANOVA. Data represents mean ± SEM of three independent healthy donors. *** p≤0.001, **** p≤0.0001.

a

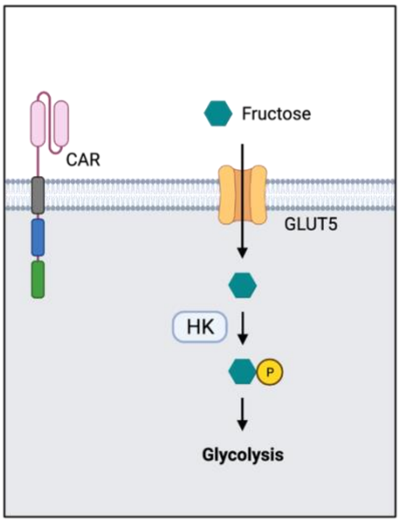


Supplementary Figure 3 | **Proposed model for fructose utilsation by CAR T cells. a.** Ectopic GLUT5 expression by CAR T cells enable trans-membrane transport of fructose into T cells, where it is phosphorylated by ubiquitously expressed hexokinase (HK) to generate the glycolytic intermediate fructose-6-phosphate (F6P). F6P is then able to enter canonical glycolysis.

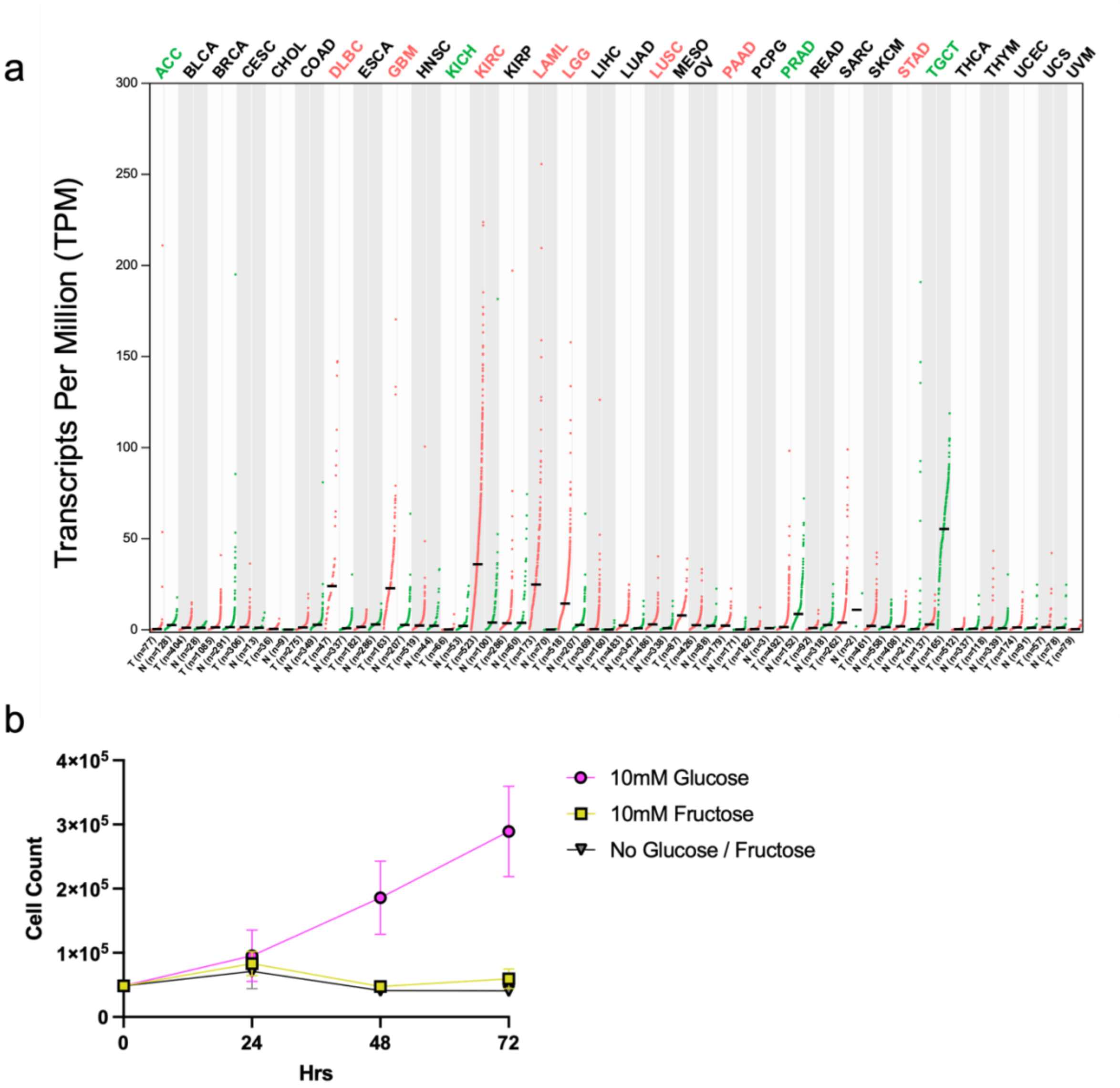


Supplementary Figure 4 | **GLUT5 expression in human cancers is heterogenous. a.** GEPIA analysis of GLUT5 transcript levels across human tumour types and matched normal tissue. Tumour types where GLUT5 expression is significantly higher than matched normal samples are colored in green and tumour types where GLUT5 expression is lower than in matched normal samples are colored in green. **b.** Growth kinetics of PC3-LN3-PSMA cells grown in media containing no glucose, 10 mM glucose or 10 mM fructose. Data represents mean ± SEM of three independent experiments.

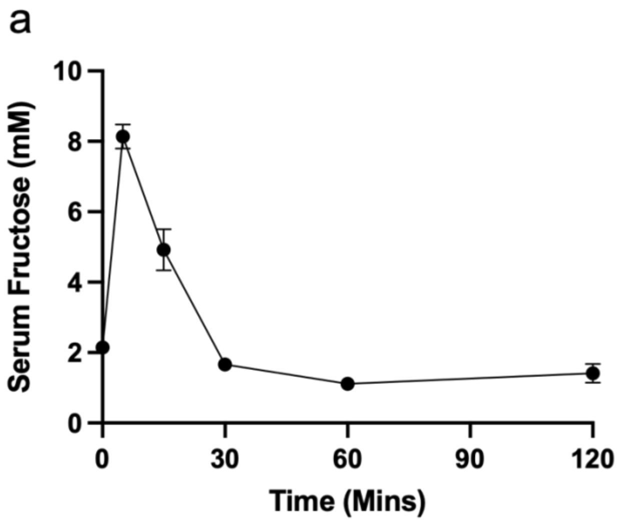


Supplementary Figure 5 | **Serum Fructose Kinetics Following Intraperitoneal (IP) Fructose Injection. a.** Fructose concentration in mice sera at defined time-points after IP injection of fructose (300 mg/kg). Data represents mean ± SEM of three independent mice.
